# Evaluation of Aggregate Oral Fluid Sampling for Early Detection of African Swine Fever Virus Infection

**DOI:** 10.1101/2025.02.19.638111

**Authors:** Bonto Faburay, Kathleen O’Hara, Marta Remmenga, Theophilus Odoom, Sherry Johnson, William Tasiame, Matilda Ayim-Akonor, Benita Anderson, Kingsley Amoake, Diane Holder, Wu Ping, Michelle Zajac, Vivian O’Donnell, Lizhe Xu, Robin Holland, Corrie Brown, Randall Levings, Suelee Robbe-Austerman

## Abstract

African swine fever (ASF) is a highly infectious viral disease that poses significant threat to the United States and global pig industries. Given the lack of effective vaccines, control and prevention of the spread of African swine fever virus (ASFV) is dependent on enhanced surveillance and early disease detection. Commercial swine operations in the US are characterized by comparatively large number of pigs, and sampling individual pigs, which represents the main strategy for current ASF surveillance, is both costly and labor intensive. The major objective of this study was to estimate the diagnostic sensitivity of pen-based aggregate oral fluid testing for ASFV in infected pigs in a pen of 30 animals and evaluate its utility as a tool to support surveillance of ASF in the United States. The study was performed in three phases: (i) Virus (Ghana ASFV24) amplification in a target host species to generate the challenge inoculum, (ii) Titration of the inoculum (10% spleen homogenate) in target host species to determine the minimum dose inducing acute ASF in pigs with survival up to 5 - 6 days post-inoculation (dpi), and (iii) The main study involving 186 pigs consisting of 6 replicates of 30 pigs per pen and one seeder pig inoculated with the Ghana ASFV24 per pen. Daily sampling of aggregate oral fluids, uncoagulated blood, oropharyngeal swabs, fecal and water nipple swabs, and recording of rectal temperatures and clinical observations, was carried out. The seeder pigs were each inoculated intramuscularly with 0.5 ml of the 10% spleen homogenate which induced the desired clinical course of ASF in the pigs with survival of up to 6 dpi. ASFV DNA could be detected in the seeder pigs as early as 1 dpi and 2 dpi in the blood and oropharyngeal swabs, respectively. Transmission of ASFV from the seeder pigs to the contact pig population was detected via positive amplification of ASFV DNA in aggregate oral fluid samples at 3 days post-contact (dpc) in 4 out of 6 pens, and in all 6 pens at 4 dpc. Testing of oropharyngeal swabs and blood samples from individual pigs revealed variable number of ASFV positive pigs between 3 and 5 dpc, with detection of 100% positivity between 6 and 18 dpc, the study endpoint. These findings demonstrate the potential utility of aggregate oral fluid sampling for sensitive and early detection of ASFV incursion into naïve swine herds. It also demonstrates that testing of environmental samples from the premises could further enhance overall ASF early detection and surveillance strategy.

**Author summary:** Early detection of ASFV in swine farms requires robust passive surveillance using sample types and sampling methods that allow sensitive and timely detection. Commercial swine operations in the US or North America are characterized by comparatively large number of pigs, and sampling individual pigs, which represents the current strategy for ASF surveillance, is both costly and labor-intensive. Oral fluid has been shown to be an acceptable sample type for detection of ASFV in individual infected pigs. For the first time on such a scale, we conducted a study enrolling 186 pigs, with daily sampling, in six experimental replicates at 3.2% pen prevalence using a highly virulent ASFV (Ghana ASFV24) to evaluate the utility of aggregate oral fluids for early detection of ASFV. Whole genome sequencing and characterization confirmed grouping of the virus with those in the p72 genotype II cluster. We demonstrate that intramuscular inoculation of the seeder pigs induces acute ASF and transmission to the contact pigs which is detectable in aggregate oral fluids as early as 3 - 4 dpc. We have shown that ASFV DNA detections in aggregate oral fluids correlate with the oropharyngeal swabs of individual pigs. In comparison, ASFV DNA in individual blood samples is detected 1-3 days later. The study demonstrates the potential utility of aggregate oral fluid sampling for enhanced surveillance of ASFV in large commercial swine operations.

## Introduction

African swine fever (ASF) is a highly contagious transboundary viral hemorrhagic disease of domestic and wild pigs presenting significant threat to the global swine industry [1–7]. Consequently, ASF is a listed disease of international concern by the World Organization for Animal Health (WOAH) [8]. It is caused by African swine fever virus (ASFV), a large double-stranded DNA virus and sole member of family *Asfaviridae* and the genus *Asfivirus*. It is a highly complex virus with a genome size that ranges from 170 to 190 kbp that encodes 150-200 proteins, including 68 structural proteins and more than 100 non-structural proteins, depending on the virus isolate [9, 10].

Originally, the emergence and outbreak of ASFV outside Africa was caused by genotype I ASFV, however, genotype II is now responsible for most of the outbreaks globally [3, 11–13]. Following its introduction in the Caucasus, Georgia, in 2007, the p72 genotype II of ASFV, has since spread globally causing disease outbreaks in Russia (2007), the Baltic States and Eastern Europe (2014), Belgium (2018), China (2018), Vietnam (2019), Cambodia (2019), North Korea (2019), Germany (2020), and most recently in the Dominican Republic (July 2021 [14–17]. The recent outbreak of ASF in the Caribbean, specifically in the Dominican Republic and Haiti, presents significant risk of introduction onto US mainland with devastating economic consequences including restrictions on international trade in swine, pork and pork products. Except for Vietnam where two vaccines have been recently approved for controlled use in the field [18–22], there are currently no effective and widely available vaccines and therefore control and prevention of spread of the virus are dependent on enhanced surveillance and early detection. Enhancing preparedness for early virus detection requires implementation of robust surveillance using sample types and sampling methods that allow sensitive and timely detection of ASFV. Commercial swine operations in the United States are characterized by comparatively large number of pigs, and sampling individual pigs for blood, the approved sample type by USDA and WOAH, which represents the main strategy for current ASF surveillance, is costly, labor intensive and unsustainable.

Oral fluid (OF) is a complex matrix containing an assortment of proteins including antibodies, mucin and a variety of enteric microorganisms as well as metabolites recovered by pigs from their environment [23, 24]. Thus, as a result of pigs’ normal exploratory behavior (smelling, tasting and biting, etc.), environmental diagnostic targets are collected in the buccal cavity [25], which may subsequently be passed onto the rope and into the oral fluid. This makes oral fluid a suitable sample type for use to support surveillance of a variety of swine pathogens [26, 27]. Oral fluid testing is based on the use of a non-invasive aggregate sampling used in industry and several laboratories in the US for diagnostic testing and surveillance for a variety of endemic swine pathogens such as swine enteric coronavirus diseases [28, 29]; porcine reproductive and respiratory syndrome [30], swine influenza A virus [31] and other viruses [32, 33]. Importantly, oral fluid has been shown to be an acceptable sample type for detection of ASFV [12, 34, 35]. Transmission of ASFV in pigs is via ingestion as the primary route, and the tonsils and oropharynx represent the sites of initial virus replication [36, 37]. Consequently, ASFV may be detected early in the pharyngeal swabs and oral fluid [38], presenting an opportunity of utilizing OF to detect viral infection prior to the onset of clinical signs. A recent simulation study concluded that following introduction of ASFV into the southeastern part of United States, a rapid response could prevent up to 79% of virus spread in the region [39]. Additionally, validation and subsequent adoption of aggregate oral fluid as an approved sample type by industry in North America and the WOAH for ASFV early detection and surveillance will significantly reduce cost of surveillance, enhance disease control, and improve profitability of the swine industry.

However, studies exploring aggregate oral fluid samples for ASFV detection have left gaps in understanding sample performance relative to pen/herd size in situations of low disease prevalence including consistency in sampling individual pigs to make conclusive interpretations and inferences. The aim of this study was to estimate the diagnostic sensitivity of pen-based aggregate oral fluid testing for ASFV in infected pigs in a pen of 30 animals and evaluate its utility as a tool to support surveillance of ASF in the United States, North America and globally.

## Materials and Methods

### Ethics statement

This study was conducted in accordance with the highest ethical standards. All experimental procedures were approved by the University of Ghana Institutional Animal Care and Use Committee (Approval number: UG-IACUC048/23-24).

### Production of ASFV inoculum (Study Phase I)

#### The virus and preparation of inoculum

A virulent ASFV isolate obtained from an ASF outbreak in a smallholder pig farm in Ghana (now named Ghana ASFV24), was used for the experimental inoculation of the pigs. The virus has been shown to be highly lethal causing 100% mortality in the affected farm. To prepare the virus inoculum for the study, a spleen tissue obtained previously from a pig that died from ASF was used. Approximately 1 gram of the tissue was transferred into a sterile mortar with addition of 0.5g of sterile sand and ground manually using a sterile pestle. Thereafter, 9 ml of sterile phosphate buffered saline (PBS) was added and the mixture further homogenized to produce a 10% tissue homogenate. Aliquots of the homogenate were transferred into 2-ml sterile microcentrifuge tubes and centrifuged at 2000 rpm for 5 min. Supernatants were transferred into clean 2-ml tubes and stored at -80 °C until used.

#### Virus amplification and determination of transmissibility

An experiment was carried out to amplify the challenge inoculum as well as assess the virulence and transmissibility of the challenge virus. Two naïve pigs weighing approximately 20 kgs each were selected for the inoculation. One pig (pig A) was inoculated intramuscularly with 2 ml of the ASFV 10% spleen homogenate. The second naïve pig (pig B) was left uninoculated and housed with the inoculated pig to facilitate virus transmission. The animals were monitored daily for onset and progression of clinical signs including pyrexia (≥ 40 °C), with daily collection and testing of whole blood. Blood samples were tested for viremia daily by quantitative PCR (qPCR) based on the protocol used at the National Veterinary Services Laboratories’ Foreign Animal Disease Diagnostic Laboratory (FADDL) [38] as described below. At a Ct value of less than 18, the inoculated pig (A) was euthanized, and spleen samples were harvested and aliquots of 10% homogenate were prepared as described above. Monitoring of the transmission and disease progression continued in pig B until it succumbed to ASF on 11dpc.

#### Whole genome sequencing and phylogenetic analysis

The ASFV inoculum sequence was analyzed by whole genome sequencing on an Oxford Nanopore Technologies (ONT. UK) GridION sequencing platform with ONT Rapid Barcoding kit (RBK114.96), and on Illumina MiSeq 2000 with Nextera XT library preparation kit and using NextSeq 2000 P3 Reagents (300 Cycles). In addition, the sample was amplified with a PCR-tiled primer set developed by Warr et al. [40], and with a p72-specific primer set [41]. The amplicons, after purification using Ampure XP beads (Beckman Coulter Life Science, US), were sequenced on ONT platform. The combined data from the different NGS runs were assembled with CLC Genomics Workbench (Qiagen, Hilden, Germany) version 24.0.2 using De Novo Assemble Long Reads and Polish with Short Reads workflow. After analyzing the contigs with the NCBI blastn tool to identify the closest isolate from the GenBank on B646L gene, the data was assembled again using the isolate as a reference in CLC. Multiple sequence alignment of generated *B646L* sequence with sequences retrieved from GenBank was performed in CLC Genomics Workbench. Maximum Likelihood phylogenetic tree was constructed in CLC with 1,000 bootstraps or iterations.

### *In vivo* dose titration (Study Phase II)

To determine the minimal dose that induces acute ASF in seeder pigs with survival of up to 5-6 dpi, three different doses of the 10% spleen homogenate, low (0.5 ml), medium (1 ml) and high (2 ml), were evaluated in 3 groups of pigs consisting of 3 animals per group. The animals were inoculated via the intramuscular route and daily whole blood samples in EDTA were collected from each animal. DNA was extracted as described below. Viral load was measured by ASFV qPCR according to the protocol described below. The animals were monitored daily for clinical signs. Clinical signs were scored according to the parameters described previously with modifications [42, 43] (S1 Table) and a clinical score of 4 or above or death served as an endpoint. The study ended on day 8 post inoculation.

### Isolation and quantification of virus inoculum

Primary swine blood macrophages were prepared from defibrinated swine blood, as described elsewhere [44]. Briefly, heparin-treated swine blood was incubated at 37C for 1-2 hours to allow sedimentation of the erythrocyte fraction. Mononuclear leukocytes were separated by a Ficoll-Paque (Pharmacia, Piscataway, NJ, USA) density gradient (specific gravity, 1.079). The monocyte/macrophage cell fraction was cultured in Primaria T-75 tissue culture flasks (Falcon, USA) containing complete macrophage media, composed of RPMI 1640 Medium (Life Technologies,) with 30% L929 supernatant, 20% heat inactivated – gamma irradiated fetal bovine serum (Sigma), 1 X antibiotics/antimycotics (Gibco), 1 X Gentamicin (Gibco) and complementary plasma for 24 hrs at 37 °C under 5% CO_2_. Adherent cells were detached from the Primaria flasks by using 10 mM EDTA in Dulbeco’s phosphate buffered saline (DPBS, Gibco), and were then reseeded into Primaria 6-well plates or 96-well plates incubated at 37 °C in 5% CO_2_, for use in assays 24 hrs later. Virus titration was performed on primary macrophage in Primaria 96-well plates. Briefly, 10-fold serial dilutions of the prepared sample (10% homogenate) were prepared in duplicate, in complete macrophage media. A volume of 50ul/ well per each dilution (4 well per dilution), by duplicate, were added to the Primary 96-well tissue culture plates seeded at a density of 5 x 10^6^ cells / plate. The plate incubated at 37 °C in 5% CO_2_. The macrophages were observed daily under the microscope up to 10 days, and detection of ASFV was determined by hemadsorption (HA). Virus concentrations are reported as HAD_50_/ml, calculated by the Reed and Muench method [45].

### Aggregate oral fluid evaluation (Phase III)

#### Animals

A total of 186 pigs, weighing 25 - 30 kgs each, were purchased from a government swine breeding facility in Accra, Ghana. The facility had no reported history of recent ASF cases or outbreak. To enroll experimental animals into the study, serum samples were collected randomly from a subset of pigs (*n* = 96) from the farm to rule out prior exposure to ASFV via detection of antibodies using a validated commercial kit according to manufacturer’s instructions (IDVet Indirect antibody ELISA, Innovative Diagnostics, Montpellier, France). The experiment to evaluate aggregate oral fluids was performed in an animal facility located at the Accra Veterinary Laboratory, Directorate of Veterinary Services and consisted of 6 pen replicates with each pen holding 30 pigs and one seeder pig (Fig. 1). Water was provided *ad libitum,* and the animals were fed twice daily.

**Fig. 1:**
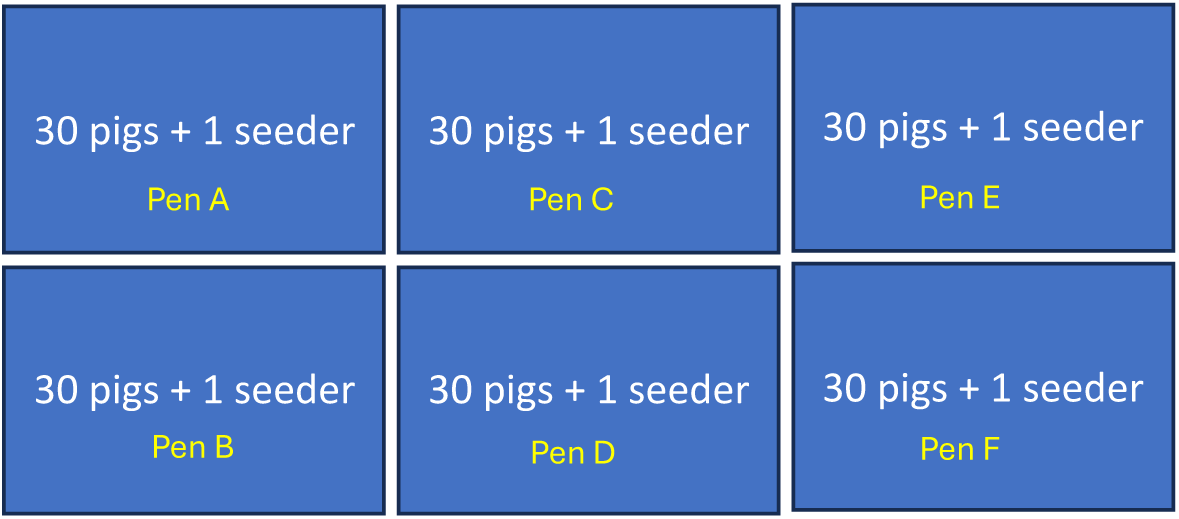
Layout of the study pens consisting of six replicates of 30 pigs and 1 seeder pig per pen. The location of the different pens (Pen A - F) is shown.

#### Inoculation of seeder pigs and contact transmission

Following acclimation and rope training, one pig from each of the 6 groups was randomly selected as “seeder pig” and inoculated intramuscular with 0.5 ml of 10% spleen homogenate of the wildtype ASFV described above. Baseline aggregate oral fluid from each pen, EDTA blood and oropharyngeal swabs from individual contact pigs were collected. Each inoculated seeder pig was introduced immediately into their respective pens to co-mingle with the contact pigs to facilitate contact transmission. This timepoint was considered 0 dpi for seeder pigs, and 0 dpc for contact pigs.

##### Sampling and clinical data collection

###### Aggregate oral fluids

Aggregate oral fluid samples were collected from contact pigs in each of the pens daily (0 - 8 dpc) and on alternate days (10 - 22 dpc). After a week of acclimation, the pigs were subjected to an initial pre-study conditioning to rope chewing for 1 hr. During each aggregate oral fluid collection, a single cotton rope (IDEXX Laboratories, Westbrook, Maine, USA) was hung in each pen on a metal crossbar at shoulder height level of the pigs. Contact pigs were allowed to chew on the rope for 30 min with monitoring of individual pig chewing activity and documentation of the proportion of pigs that chewed on the rope on any given day. The rope from each pen was collected and oral fluid was squeezed out into a tube and transported to the laboratory for further processing. During aggregate oral fluid collection, seeder pigs were isolated in specifically designed pens to prevent them from chewing on the rope and were released back into their respective pen population after completion of each sampling to facilitate contact transmission.

###### Blood and oropharyngeal swabs

Uncoagulated blood samples in ethylenediaminetetraacetic acid (EDTA) in vacutainer tubes were collected daily from individual pigs from day 0 to day 8 dpc, and on alternate days until 22 dpc, the study endpoint. The samples were temporarily stored at -80 °C. Prior to use, the samples were centrifuged at 3,000 rpm for 30 min and the supernatant collected for DNA extraction as described above. Oropharyngeal blood swabs were also taken daily using individual sterile polyester tipped swabs in 1 ml of viral transport medium (VTM) (Teknova, Hollister, CA) daily from individual pigs from 0 to 8 dpc, and on alternate days 10 - 22 dpc, until the study endpoint. The samples were stored at -80 °C before processing or processed for DNA extraction immediately.

###### Environmental samples: fecal and water nipple swabs

To evaluate potential utility of environmental sampling to support ASF surveillance, fecal swabs were collected daily from the middle and the four corners of each pen using sterile polyester tipped swabs in 1 ml of sterile PBS from 0 to 8 dpc. Similarly, water nipple swabs were also taken from each pen using individual sterile polyester tipped swabs in 1 ml of VTM from 0 to 10 dpc. These samples were stored at -80 °C until processed for DNA extraction.

###### Clinical data

Rectal temperatures and clinical scores [42, 46] of individual pigs based on previously described the criteria (S1 Table) were recorded daily until the end of the study on 22 dpc. In cases where death did not occur overnight, pigs demonstrating a clinical score equal or greater than 4 were euthanized. Necropsies were performed on a subset of pigs to collect representative samples and ascertain the cause of death.

### DNA extraction and quantitative real-time PCR (qPCR)

DNA extraction from aggregate oral fluids, whole blood, oropharyngeal swabs, water nipple and fecal swabs was performed using Qiagen DNA extraction kit according to manufacturer’s instructions with modifications using 100 µl of sample added to 120 µl of lysis buffer, 20 µl proteinase K and 3 µl internal control Xeno. The DNA was eluted in 50 µl elution buffer. ASFV genomic DNA was detected using quantitative real-time PCR targeting the p72 open reading frame [38]. Beta actin was used as the internal extraction and amplification control [47]. The qPCR reaction was carried out as described previously [34] in a reaction mix containing Taqman Fast 1-step master mix (ThermoFisher Scientific, Inc.) and was amplified using the Bio-Rad CFX96 instrument (Bio-Rad Laboratories Canada Ltd, Missisauga, Ontario) in 45 amplification cycles (50 °C for 5 minutes, 95 °C for 20 seconds, 95 °C for 3 seconds and 60 °C for 30 seconds).

### Data analysis

For comparing sensitivity of ASFV DNA detection in aggregate oral fluids with detection in individual blood and oropharyngeal swabs collected from contact pigs, we derived the proportion of pens and pigs (for blood and oropharyngeal swabs) positive at each time point post contact.

For blood and oropharyngeal swabs, due to the large variations in the percentage of positive animals per time point, we derived the median positivity for each time point. To analyze the trend in Ct values (viral load) for contact pigs across pens, we derived mean pen Ct values with standard deviations at each timepoint. These values were plotted to demonstrate the trend over time in the Ct value changes, indicative of changes in viral load. Kaplan Meier survival estimate was used to analyze time to death and survival rates for the dose titration study. Data was analyzed using GraphPad Prism 10.

## Results

### Production and amplification of virus inoculum

Inoculation of Pig A with 10% spleen homogenate with the virulent wildtype ASFV resulted in acute ASF exhibiting characteristic clinical signs including elevated rectal temperature, cutaneous cyanosis, depression and lethargy. Upon necropsy, the animal exhibited typical for ASF gross pathological changes and lesions such as splenomegaly, hemorrhage and lymphadenopathy. The animal developed detectable viremia as determined by qPCR at 1 dpi with progressive increase in viral load as demonstrated by reduction in cycle threshold (Ct) values from Ct 35 on 1 dpi to Ct 15 on 4 dpi. The pig was euthanized on 4 dpi, and the spleen was harvested and stored at -80 °C and later used to prepare 10% homogenate for use as challenge inoculum in the *in vivo* dose titration and in subsequent experiments. Pig B that was co-housed with Pig A developed detectable viremia on 8 dpc as well as clinical symptoms characteristic of acute ASF thus confirming transmissibility of the virus from infected to naïve hosts. Pig B died from ASF on 11 dpc.

### In vivo dose titration

To determine the minimum dose that would induce acute ASF in the seeder pigs with survival of up to 5 - 6 dpi, three different doses of the 10% spleen homogenate, low (0.5 ml), medium (1 ml) and high (2 ml), were evaluated as described above in 3 groups of pigs, group A, B and C, respectively. In vitro quantification of the virus inoculum in porcine primary macrophages indicated a titer of 4.43 HAD_50_/ml.

Animals in all three groups exhibited elevated temperatures (data not shown) and developed viremia at 1 dpi (Ct value range: 22.04 - 33.34). There was progressive reduction in Ct values with increasing severity clinical symptoms among animals in all three groups, Group A (Ct value range: 14.1 - 16.1), Group B (Ct value range: 15.96 -16.71) and Group C (Ct value range: 14.97 - 15.8). Comparing responses among the different groups, animals that received the 0.5 ml homogenate (4.43 HAD_50_/ml) in Group A exhibited the longest survival time of 6 days without a mortality event (Fig. 2). Consequently, this dose was used to inoculate the seeder pigs in the aggregate oral fluid evaluation study.

**Fig. 2.**
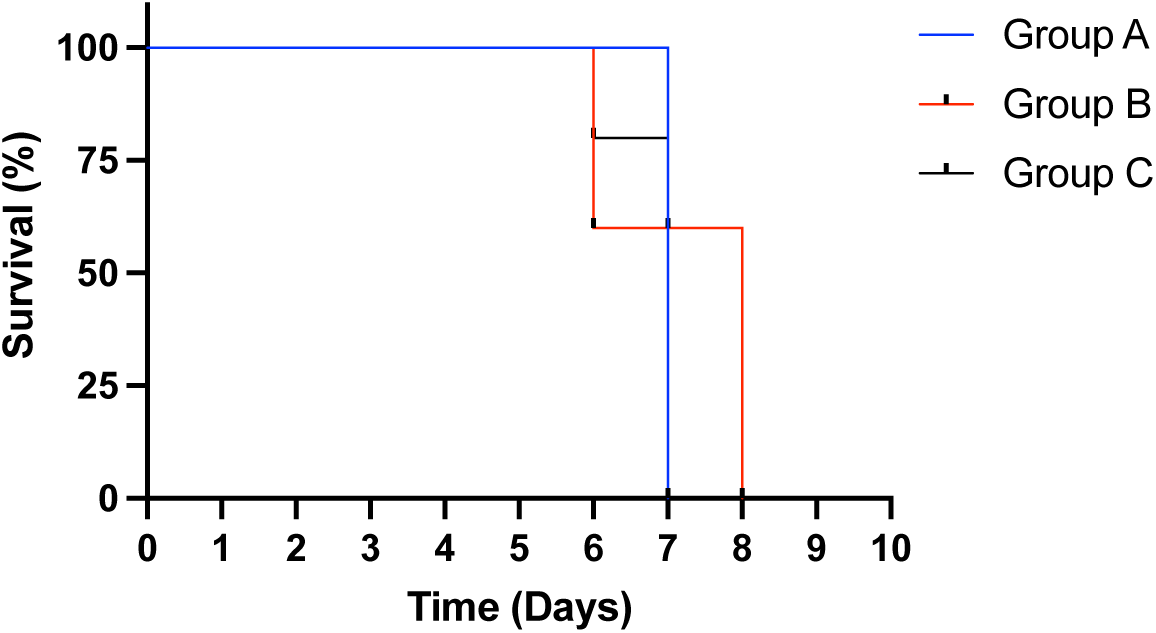
Kaplan Meier survival analysis showing time of survival for animals in the different dose groups. Only pigs in Group A (0.5 ml) survived to 6 dpi without a mortality event.

### Whole genome sequencing and phylogenetic analysis

Whole genome sequencing of ASFV inoculum revealed a genome size of approximately 185 kbp which was within the size-range of ASFV genome (170 - 190 kbp). Phylogenetic analysis based on the *B646L* gene sequences (encoding the p72 capsid protein) showed the virus inoculum grouped in the genotype II cluster together with ASFV Georgia 2007 isolate (S1 Fig.). The comparison at the p72 nucleotide sequence level revealed the virus inoculum was 100% identical to the previously published sequences of an ASFV isolated in Ghana [48].

### Aggregate oral fluid evaluation

#### Virus inoculation and clinical observations

Inoculation of the seeder pigs with 0.5 ml of spleen homogenate resulted in elevated rectal temperature response by 2 - 3 dpi and onset and development of clinical signs by 2 - 4 days (Fig. 3A1 and 4A). The majority of contact pigs in the six pens exhibited increase in rectal temperatures starting 10 - 11 dpc, and in several cases a decrease in rectal temperature below the normal in pigs exhibiting severe clinical signs or in the terminal stages of disease progression (Fig. 3A-F). Clinical observations in contact pigs depicted an onset of rising mean clinical scores on 7 - 8 dpc in pen A, B, E and F, and on 10 dpc in pen C and D (Fig. 4B).

**Fig. 3.**
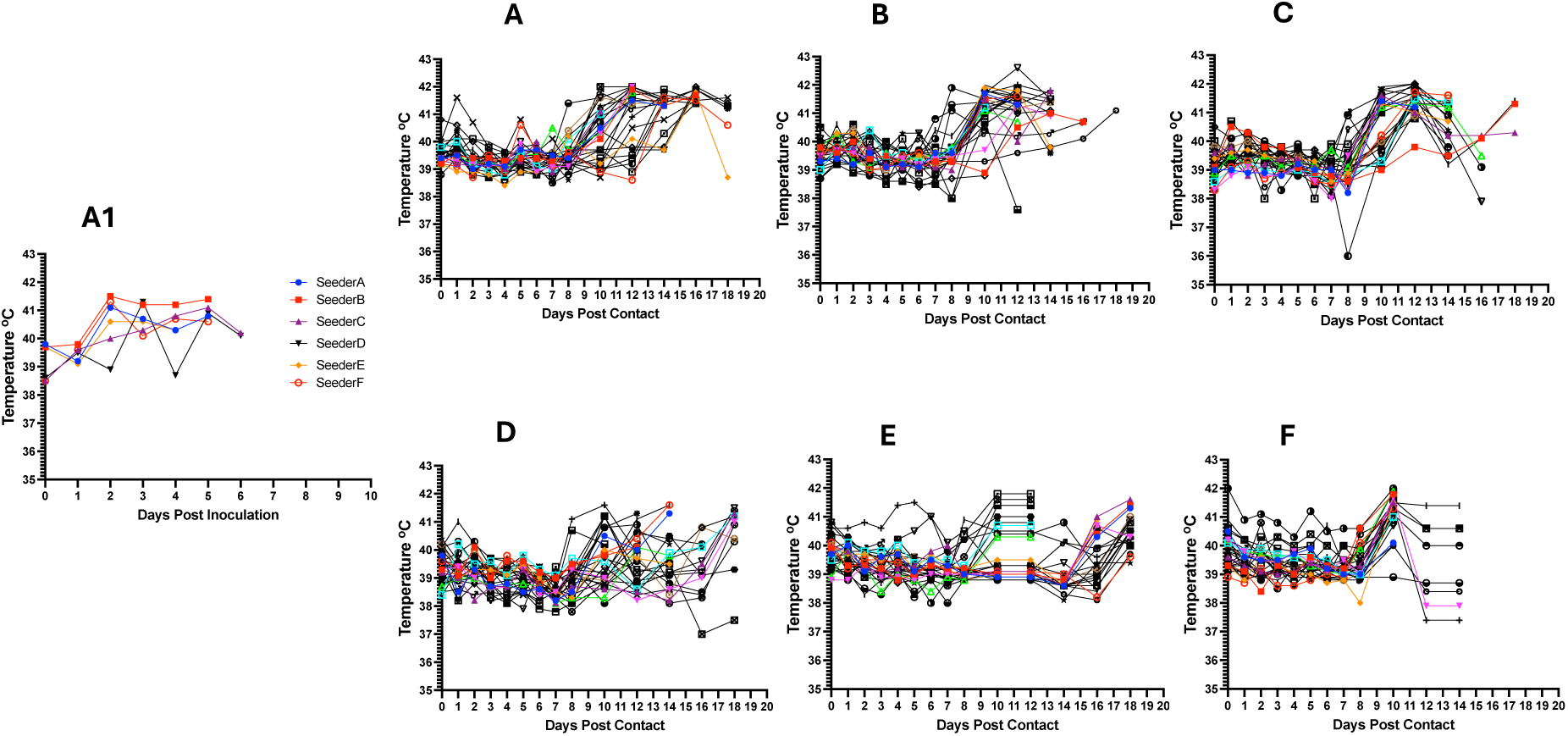
Kinetics of rectal temperature in contact pigs in response to ASFV infection. The overall trend shows increasing rectal temperatures from 10 dpc onwards with the onset of severe clinical symptoms (A-F); each temperature curve represents response of an individual pig. Temperature response in seeder pigs shows increase on 2-3 dpi (A1).

**Fig. 4.**
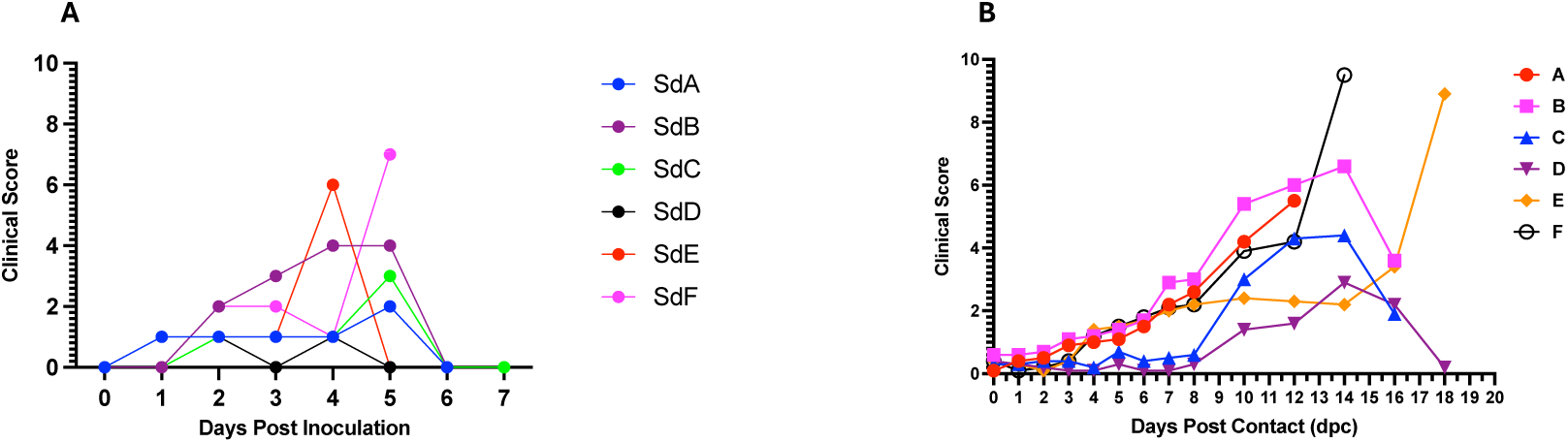
Dynamics of clinical scores in seeder (A) and in contact pigs (mean score) for each per pen (B). The data show rising clinical scores over time with onset of clinical signs and progression of the disease.

### ASFV genomic detection in seeder pigs

Blood samples collected from all six seeder pigs tested positive for ASFV DNA on 1 dpi, with Ct values ranging from 30.82 to 38.66 (Fig. 5A, Table 1). Ct values for all seeder pigs declined with the onset of clinical signs and disease progression from mean Ct value of 35.75 on 1 dpi to 16.4 on 5 dpi, and 15.8 on 6 dpi. In contrast, ASFV DNA was detectable in the oropharyngeal swabs at a later time-point on 2 dpi (Ct value range: 27.6 - 36.2) (Fig. 5B, Table 1). Ct values for all seeder pigs declined temporally with the onset of clinical signs and disease progression from mean Ct value of 32.64 on 2 dpi to 22.48 on 5 dpi, and 22.3 on 6 dpi.

**Fig. 5.**
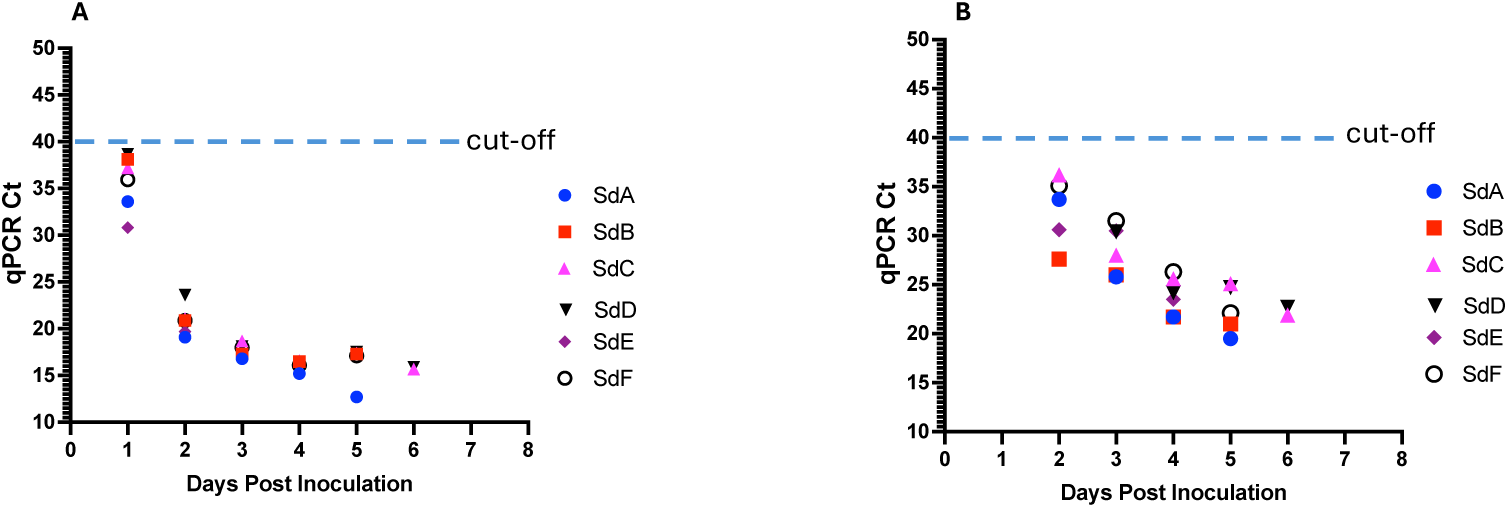
Kinetics of ASFV DNA in seeder pigs in response to intramuscular inoculation with 0.5ml of 10% spleen homogenate. Detection of DNA in blood (A) and in oropharyngeal swabs (B).

**Table 1.** Summary of mortality and ASFV genomic detection in aggregate oral fluid, blood and oropharyngeal swabs of seeder and contact pigs.

| Pen ID | Seeder pig mortality (dpi) | Initial positive OF detection (dpc) | Initial positive detection in seeder pigs (dpi) |  | Initial positive detection in contact pigs (dpc) |  |
| --- | --- | --- | --- | --- | --- | --- |
|  |  |  | Blood | Oph swab | Blood | Oph swab |
| A | 6 | 3 (35.46) | 1 (33.6) | 2 (33.7) | 6 (34.34 - 37.4) | 3 (35.21 - 37.10) |
| B | 6 | 3 (35.63) | 1(38.12) | 2 (27.6) | 5 (37.19) | 3 (35.95 - 39.83) |
| C | 7 | 3 (35.85) | 1(37.21) | 2 (36.2) | 5 (36.20 - 39.41) | 3 (37.51 - 38.50) |
| D | 7 | 4 (35.7) | 1 (38.66) | 3 (30.4) | 5 (32.67 - 33.40) | 4 (34.11 - 39.04) |
| E | 5 | 3 (36.14) | 1 (30.82) | 2 (30.6) | 5 (32.78 - 36.78) | 3 (35.19 - 38.62) |
| F | 5 | 4 (35.68) | 1 (35.94) | 2 (35.01) | 5 (35.76 - 38.41) | 4 (35.61 - 37.71) |

Note: qPCR Ct values are shown in parenthesis; OF = oral fluid; Oph = oropharyngeal

### ASFV genomic detection in blood and oropharyngeal swabs of contact pigs

To determine the infection status and kinetics of ASFV transmission among contact pigs and how these correlated with aggregate oral fluid detections at the pen level (Pen A-F), we tested blood and oropharyngeal swabs of individual pigs from 0 through 18 dpc, the end of the sampling period.

#### Detection in Pen A

Initial detection of ASFV genomic material in the blood of contact pigs occurred on 6 dpc in 33.3% (10/30) of pigs (Ct value range: 38.32 to 34.84) (Fig. 6A1; Table 1). The proportion of pigs that tested positive increased to 50% on 7 dpc (Ct value range: 28.36 - 38.56) and to 100% on 8 dpc (Ct value range: 39.32 to 17.95) (Fig. 7A). There was a decreasing trend in Ct values from 1 dpc through 18 dpc (Fig. 6A1). Seeder pig A died on 6 dpi. For oropharyngeal swabs, initial detection of ASFV genomic material occurred on 3 dpc in 56.7% (17/30) of pigs (Ct value range: 32.21-37.4) (Fig. 6A2; Table 1). The proportion of pigs testing positive increased to 96.6% (29/30) on 5 dpc (Ct value range: 30.12 - 37.91) and 100% on 7 dpc (Ct value range: 24.13 - 38.19) (Fig. 7B). Similarly, there was a decreasing trend in Ct values as described above (Fig. 6A2).

**Fig. 6.**
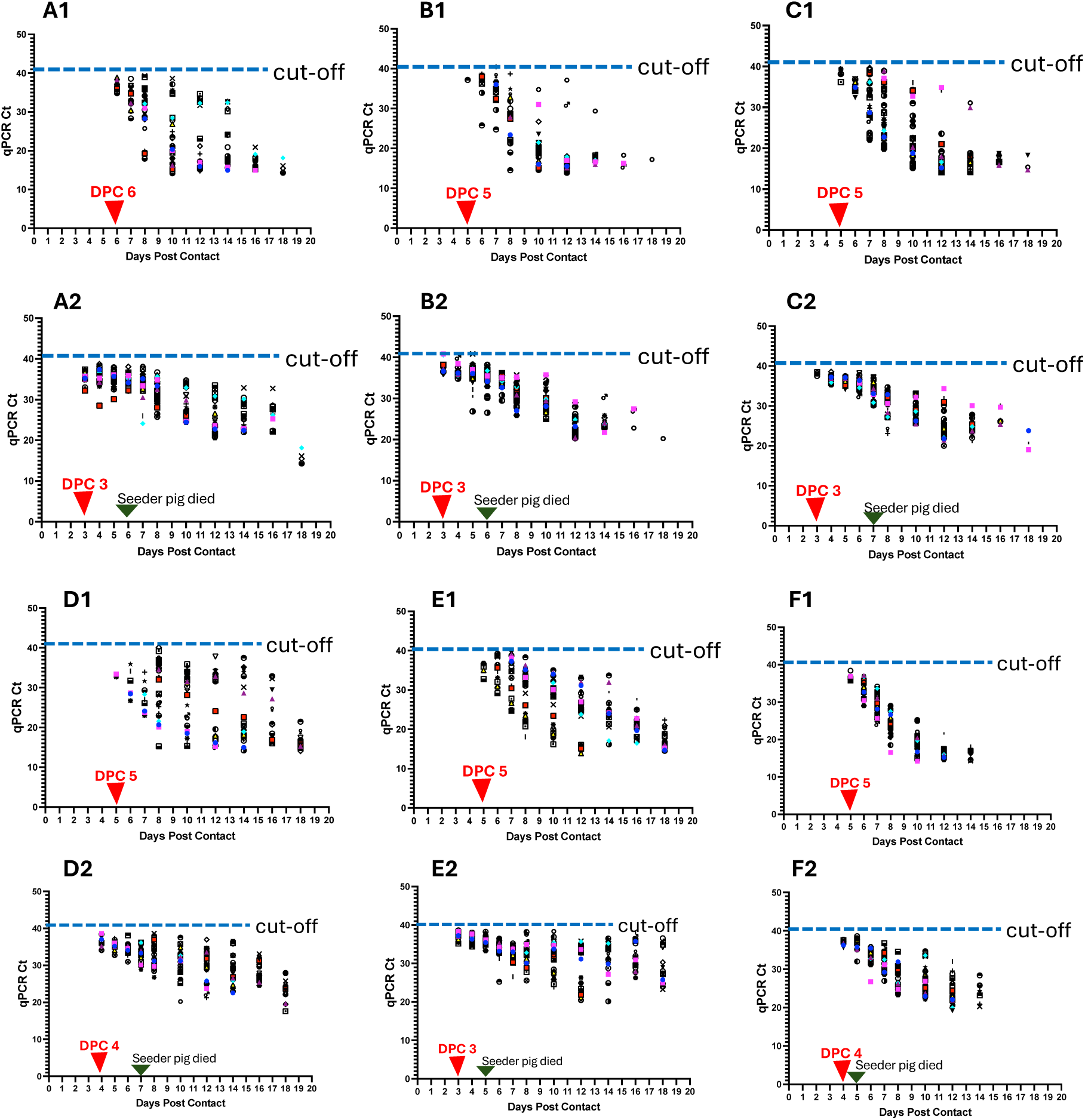
ASFV genomic detection in individual pigs in the six pens (A-F). Detection of ASFV genomic material in the blood of individual pigs per pen (A1 – F1) and in the oropharyngeal swabs (A2 - F2). Data show earlier detection of ASFV DNA in oropharyngeal swabs than in blood. The seeder pigs died 5-7 days post inoculation.

**Fig. 7.**
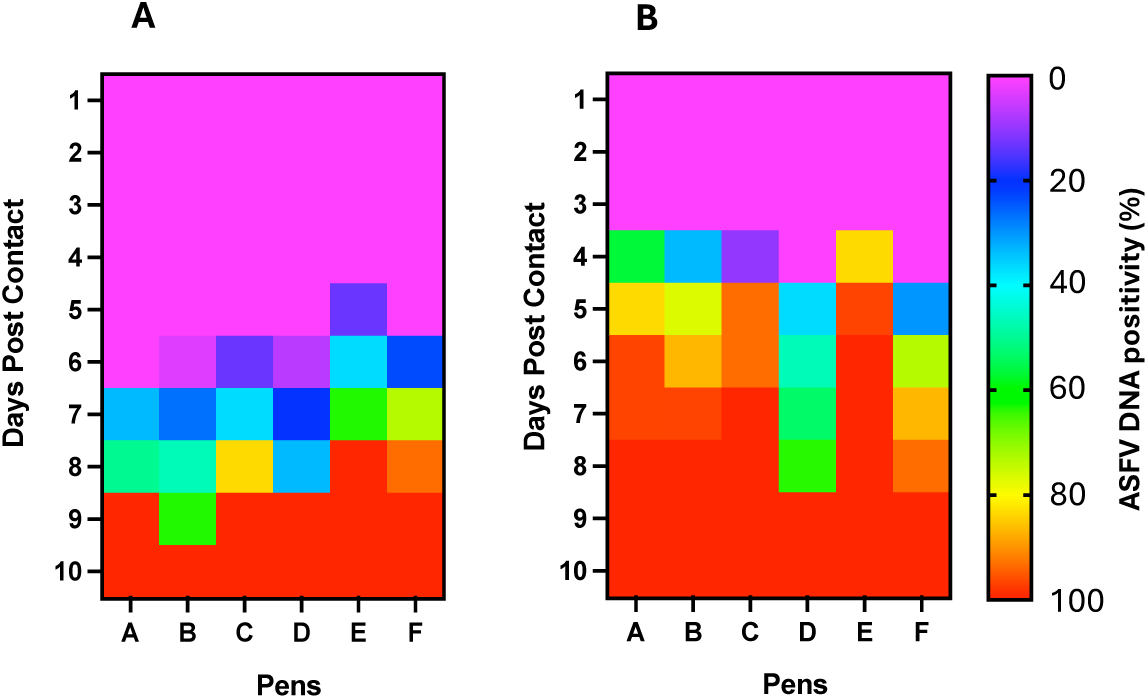
Heat map of ASFV DNA positivity in blood (A) and oropharyngeal swabs (B). Both figures display temporal increase in ASFV positivity and also demonstrate earlier detection of ASFV DNA in the oropharyngeal swabs than in blood.

#### Detection in Pen B

Initial detection of ASFV genomic material in the blood of contact pigs occurred on 5 dpc in a pig indicating 3.3% positivity (Ct value: 37.19) (Fig. 6B1; Table 1). The proportion of pigs that tested positive increased to 26.7% (8/30) on 6 dpc (Ct value range: 25.79 - 39.12) and 100% positivity on 10 dpc (Ct value range: 14.60 - 34.77) (Fig. 7A). There was a decreasing trend in Ct values from 1 dpc through18 dpc (Fig. 6B1). Seeder pig B died on 6 dpi. For oropharyngeal swabs, initial detection of ASFV genomic material occurred on 3 dpc in 10 pigs (33% positivity) within a Ct value range of 35.95 - 39.83 (Fig. 6B2; Table 1). The proportion of positive animals increased to 76.7 (23/30) on 4 dpc (Ct value range: 34.82 - 39.92), 86.7% (26/30) on 5 dpc (Ct value range: 26.83 - 39.83) and 100% on 7 dpc (Ct value range: 29.50 - 35.51) (Fig. 7B). There was a decreasing trend in Ct values over time as described above (Fig. 6B2).

#### Detection in Pen C

Initial detection of ASFV DNA in the blood of contact pigs occurred on 5 dpc in 4 pigs (13.3% positivity) within a Ct value range of 36.2 - 39.41 (Fig. 6C1; Table 1). Thirty seven percent (11/30) of the pigs tested positive on 6 dpc (Ct value range: 32.43 to 37), 83.3% (25/30) on 7 dpc (Ct value range: 28.17 - 39.61) and 100% on 8 dpc (Ct value range: 20.28 to 38.89) (Fig. 7A). There was a decreasing trend in Ct values from 1 dpc through 18 dpc (Fig. 6C1). Seeder pig C died on 7 dpi. For oropharyngeal swabs, initial detection of ASFV DNA occurred on 3 dpc in 3 pigs (10% positivity) with Ct values 38.5, 37.51 and 38.3 (Fig. 6C2; Table 1). The proportion of pigs that tested positive increased to 93.3% (28/30) on 4 dpc (Ct value range: 35.25 - 38.57). There was 100% positivity on 6 dpc (Ct value range: 32.45 - 38.21) (Fig. 7B). Similarly, there was a decreasing trend in Ct values over time (Fig. 6C2).

#### Detection in Pen D

Initial detection of ASFV DNA in the blood of contact pigs occurred on 5 dpc in 2 pigs (6.7% positivity) within a Ct value range of 32.67 - 33.4 (Fig. 6D1; Table 1). The proportion of pigs that tested positive increased on 6 dpc to 20% (6/30) (Ct value range: 32.43 - 37) and on 7 dpc to 33.3% (10/30) (Ct value range: 26.71 - 35.96). There was 100% positivity on 8 dpc (Ct value range: 15.24 - 39.19) (Fig. 7A) and a decreasing trend in Ct values from 1 dpc through 18 dpc (Fig. 6D1). Seeder pig D died on 7 dpi. For oropharyngeal swabs, initial detection of ASFV DNA occurred on 4 dpc in 11 pigs (36.7% positivity) within a Ct value range of 34.11 - 39.04 (Fig. 6D2; Table 1). The proportion of pigs that tested positive increased to 46.7% (14/30) on 5 dpc (Ct value range: 34.11 - 37.95), 53.3% (16/30) on 6 dpc (Ct value range: 30.99 - 38.10), 63.6% (21/30) on 7 dpc (Ct value range: 28.89 - 36.22) and a100% positivity on 8 dpc (Ct value range: 32.45 - 38.21) (Fig. 7B). There was a decreasing trend in Ct values as described above (Fig. 6D2).

#### Detection in Pen E

Initial detection of ASFV DNA in the blood of contact pigs occurred on 5 dpc in 4 pigs (13.3% positivity) within a Ct value range of 32.78 - 36.78 (Fig. 6E1; Table 1). Eleven pigs (36.7%) tested positive on 5 dpc (Ct value range: 29.19 - 39.22) and 19 pigs (63.3%) on 7 dpc (Ct value range: 24.72 - 39.46). All the pigs (100%) tested positive on 8 dpc (Ct value range: 18.07 - 38.32) (Fig. 7A). There was a time-dependent decrease in Ct values on 1 dpc through 18 dpc (Figure 6E). Seeder pig E died on 5 dpi. For oropharyngeal swabs, initial detection of ASFV DNA occurred on 3 dpc in 25 pigs (83.3% positivity) within a Ct value range of 35.19 - 38.62 (Figure 6E2; Table 1). The proportion of pigs that tested positive increased to 96.7% (29/30) on 4 dpc (Ct value range: 35.21 - 38.2). There was 100% positivity on 5 dpc (Ct value range: 33.32 - 38.27) through to 10 dpc (Fig. 7B) and a temporal reduction in Ct values from 1 dpc through 18 dpc (Fig. 6E2).

#### Detection in Pen F

Initial detection of ASFV DNA in the blood of contact pigs occurred on 5 dpc in 7 pigs (23.3% positivity) within a Ct value range of 35.56 - 38.41 (Fig. 6F1; Table 1). Twenty-two animals (73.3%) tested positive on 6 dpc (Ct value range: 30.45 - 37.12) and 28 pigs (93.3%) on 7 dpc (Ct values range: 24.13 - 33.64). There was 100% positivity on 8 dpc (Ct values range 14.22 - 27.52) and afterwards (Fig. 7A). The Ct values decreased over time from 1 dpc through 14 dpc (Fig. 6F1). Seeder pig F died on 5 dpi. Initial detection of ASFV DNA in the oropharyngeal swabs occurred on 4 dpc in 9 pigs (30% positivity) within a Ct value range of 32.04 - 35.6 (Fig. 5F2; Table 1). The proportion of pigs testing positive increased to 73.3% (22/30) on 5 dpc (Ct value range: 34.11 - 35.97), 86.7% (26/30) on 6 dpc (Ct value range: 26.74 - 35.97) and 100% on 8 dpc (Ct value range 24.63 - 30.89) and afterwards (Fig. 7B). There was a temporal reduction in Ct values from 1 dpc through 14 dpc (Fig. 6F2).

### ASFV genomic detection in aggregate oral fluids

To evaluate the sensitivity of detecting ASFV genomic material in aggregate oral fluids, we examined detection performance at an ASF pen prevalence of 3.2%. Across the pens, the average proportion of pigs that chewed on the rope was above 60% (range 62 - 79%) from 1 to 10 dpc. From 11 dpc and onwards, the numbers declined progressively with the onset and severity of clinical signs (Fig. 8A).

**Fig. 8.**
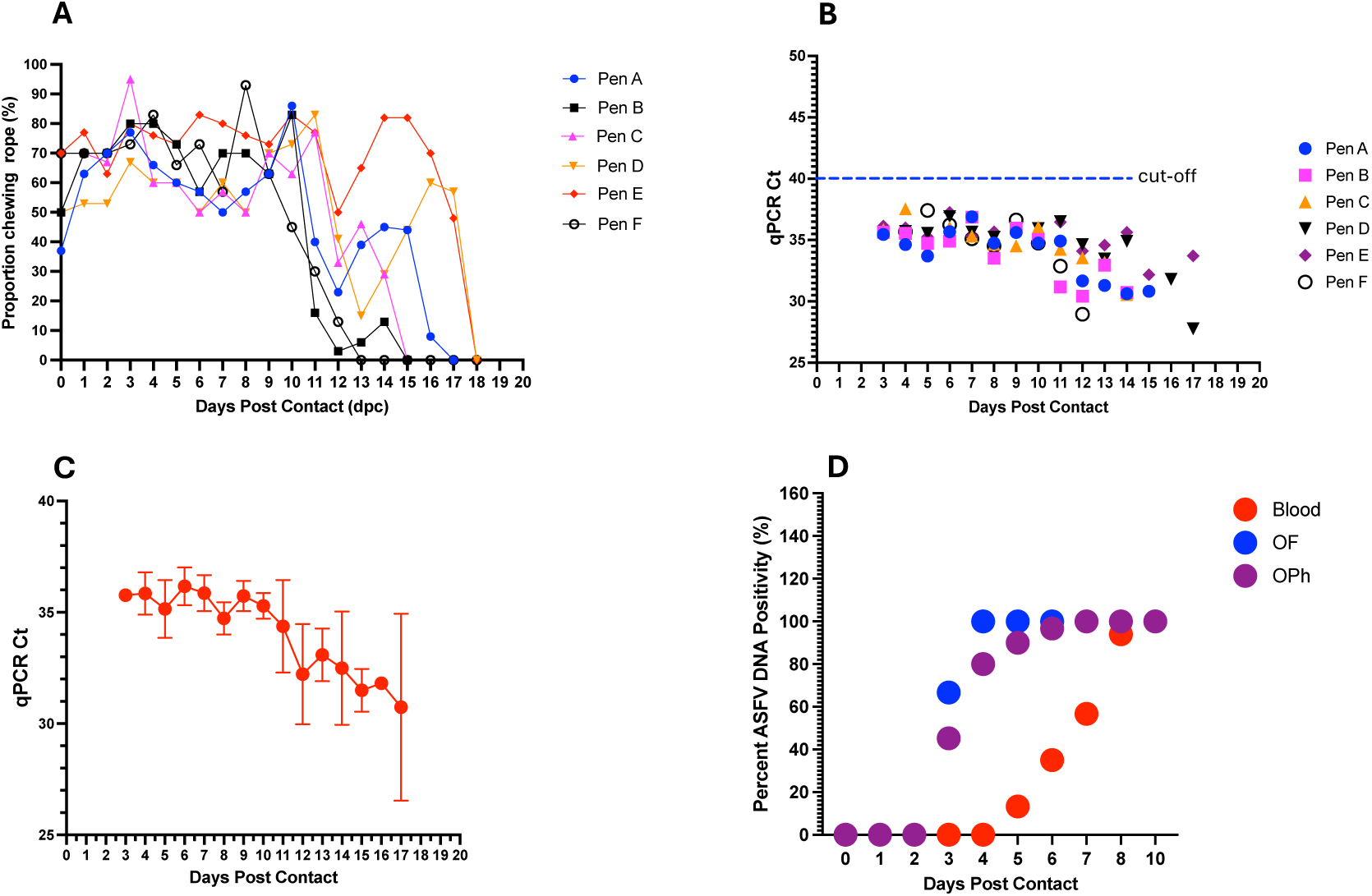
Evaluation of aggregate oral fluids. Dynamics in the proportion of pigs chewing on the rope in the different pens (A-F) at different timepoints post contact. On average more than 60% of pigs in each pen chewed on the rope from 1 - 10 dpc (A). ASFV DNA detection in aggregate oral fluids with initial positive detections on 3 dpc in 4 of the 6 pens, and in all pens on 4 dpc and afterwards (B). Genomic detection at different time-points demonstrates decreasing trend in mean Ct values with disease progression and increase in viral load (C). Comparison of the sensitivity of ASFV DNA detection in whole blood, oropharyngeal (OPh) swabs and aggregate oral fluids (OF). Detection of ASFV DNA occurred earlier in aggregate oral fluid and oropharyngeal swabs (on 3 dpc) than in blood (5 dpc) of contact pigs via contact transmission.

Initial detection of ASFV DNA in aggregate oral fluids occurred on 3 dpc in 4 out of the 6 pens, Pen A (Ct value 35.46), Pen B (Ct value 35.63), Pen C (Ct value 35.85) and Pen E (Ct value 36.14) (Fig. 8B and S2 Table). Aggregate oral fluids in all six pens (100% pen detection rate), including Pen D and Pen F occurred on 4 dpc within a Ct value range of 34.63 - 37.55 (Fig. 8B and S2 Table). Thereafter, ASFV DNA was consistently detected in all pens (A-E) until 14 dpc after which the pigs became too depressed and lethargic to chew on the rope (Fig. 8A and S2 Table). Overall, there was a decreasing trend in the average Ct values with disease progression, with mean Ct values decreasing from 35.85 on 4 dpc to 30.74 on 17 dpc (Fig. 8C).

Comparing the sensitivity of ASFV DNA detection in aggregate OF with that in blood and oropharyngeal swabs, viral DNA was detected in aggregate oral fluid as early as 3 dpc (Ct value range: 35.46 - 36.14) (Fig. 8D; Table 1), in contrast to 5 dpc (Ct value range: 32.67 - 39.41) in blood (Fig. 8D; Table 1). Detections in oropharyngeal swabs of individual pigs correlated with aggregate OF detections with initial detection in the former sample type also occurring on 3 dpc and afterwards (Fig. 8D). Aggregate OF from 4 out of 6 pens (67%) tested positive on 3 dpc, and 100% on 4 dpc (Ct value range: 34.63 - 37.55) and afterwards until 10 dpc (Fig. 8D). Initial detection rate in oropharyngeal swabs on 3 dpc was comparatively lower showing a median pen positivity of 45.2% (Fig. 8D). For detection in individual blood samples, the earliest time-point of positive DNA detection was 5 dpc with a median pen positivity of 13.3%, followed by a gradual increase in positivity at subsequent time-points to 35% (6 dpc), 56.7% (7 dpc), 94% (8 dpc) and 100% (10 dpc) (Fig. 8D).

### Environmental sample testing: water nipple and fecal swabs

In a preliminary evaluation of environmental samples, water nipples and fecal swabs, to support ASF surveillance, we performed daily pen-level sampling via from 0 - 10 dpc. The qPCR results exhibited variability in detection performance amongst individual pens at different time-points. For water nipples swabs, there was a positive ASFV DNA detection in Pen E, on 2 dpc (Ct value 36.3). Four pens (A, B, C and D) out of 6 were positive on 4 dpc (Ct values 34.4, 35.1, 36.9 and 35.5), a 67% pen detection rate (S3 Table). Thereafter, there were inconsistent and sporadic positive DNA detections until 9 dpc (S3 Table). The earliest positive ASFV DNA detection in fecal samples occurred on 4 dpc in three Pens (A, B and D) (Ct values: 31.33, 35.88 and 33.88) indicating a 50% pen-level detection rate (S3 Table). Thereafter, there was inconsistent and sporadic detections in the various pens until 8 dpc. Four pens (Pen A, D, E and F) out of 6 tested positive (Ct values 33.45, 35.75, 34.12 and 32.9) indicating a 67% pen-level detection rate. All six pens (A - F) tested positive (Ct values 29.89, 35.47, 31.7, 34.04, 34.09 and 31.17) on 10 dpc, a 100% detection rate (S4 Table).

### Mortality in contact pigs

Mortalities among contact pigs occurred between 10 and 22 dpc with the earliest deaths occurring in Pen A and B on 10 dpc (Fig. 9). The first deaths in Pen C, D, E and F occurred on 12 dpc (Fig. 9). There was non-uniformity in the mortality curves for the different pens with Pen F showing the highest number of daily mortalities occurring between 12 and 16 dpc (Fig. 9F). Cumulatively, the highest number of deaths on a single day across all six pens was observed between 12 and 15 dpc accounting for more than half (61%, 110/180) of the mortalities (Fig. 9G).

**Fig. 9.**
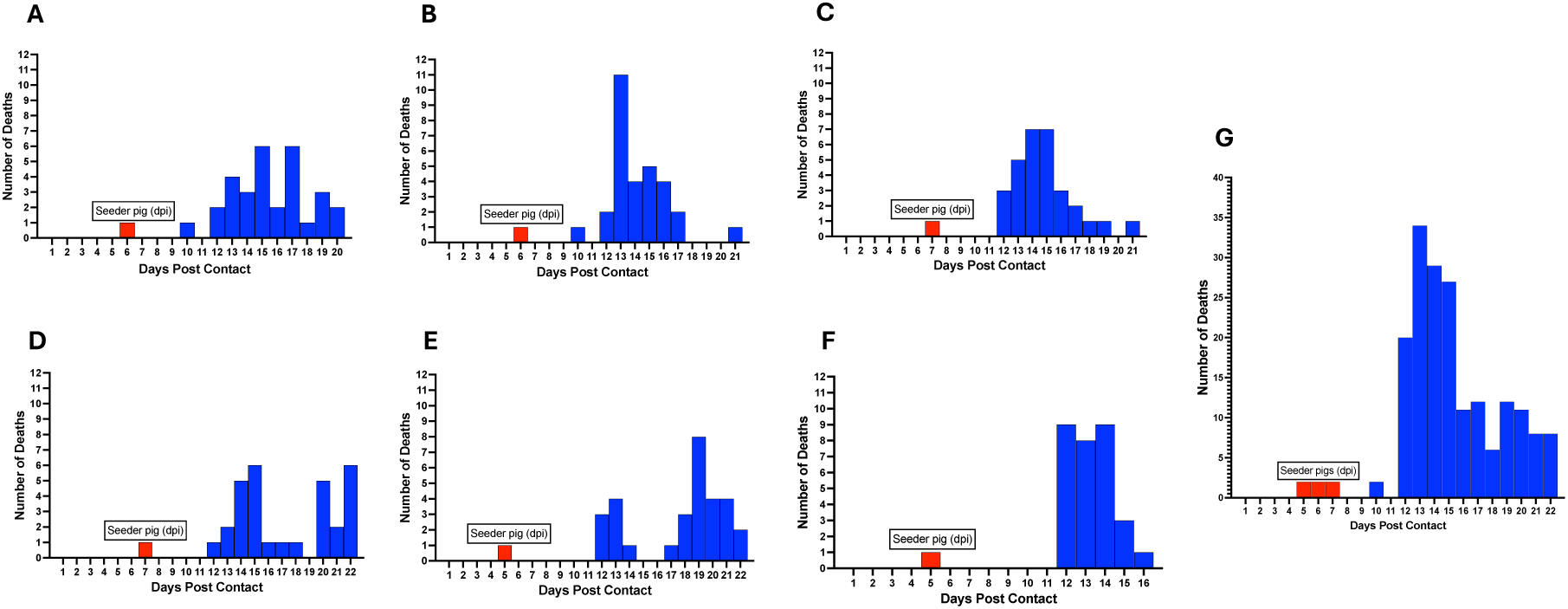
ASF mortality curve in individual pens (A-F) showing variability in the death rate at different time-points post contact. Mortalities occurred between 10 - 22 dpc. The cumulative mortality curve for all six pen pens is shown (G). Red bars denote day of seeder pig death post inoculation and introduction in each pen.

## Discussion

African swine fever is a highly contagious viral disease of domestic and wild pigs that poses significant threat to the global swine industry [6, 49, 50]. There is no effective and widely available vaccine, and therefore prevention in the US and North America generally is dependent on surveillance and early detection. Current surveillance strategy relies on collection of individual blood samples, the approved sample type for live pigs, for early ASF detection. This approach is laborious and unsustainable in commercial operations with large number of pigs. Our study represents the first of its kind that systematically and consistently collected daily aggregate oral fluids as well as individual blood and oropharyngeal swabs from contact pigs across six pen replicates involving 186 pigs to evaluate the use of aggregate oral fluid as an alternative sampling method for ASF surveillance in US and North America. Unlike previous studies, we performed initial amplification of the virus inoculum in the target host species in phase 1 of the study to confirm virulence, minimize possible attenuating mutations that may result from *in vitro* cell culture propagation, and to ensure the virus retained transmissibility to contact hosts. Additionally, we performed an *in vivo* pre-study, in phase II, to determine the minimum dose of the virus that induces acute ASF in the target host with survival up to 5 - 6 days to generate the seeder pigs.

Several studies have been carried out previously that evaluate the utility of aggregate oral fluid for early detection and surveillance of ASFV [12, 34, 35]. These studies vary in experimental design as well as ASFV pen prevalence, virus phenotype, sample size, including sampling strategy. In one study using moderately virulent ASFV Malta’78 (genotype I) strain [51] and a 50% pen prevalence (n = 10) and intranasal inoculation of seeder pigs (n = 5), ASFV genome has been detected in the oropharyngeal swabs of the contact pigs around 4 dpc regardless of dosage [46]. In other studies, using ASFV Georgia 2007/1 (genotype II) and ASFV Lisbon 60 (genotype I) strains administered via intramuscular route and based on small sizes with ASFV pen prevalence of 50 - 100%, ASFV genomic material has been detected in the aggregate oral fluids on 2-3 dpi [35, 52]. These studies unlikely represent the most probable scenario of ASFV introduction into naïve swine herds whereby 50% of the herd get infected simultaneously with equal risk of exposure to ASFV and subsequent early detection via surveillance.

Recently, a study has been performed using highly virulent Georgia 2007/1, and a moderately virulent Malta’78, strain. The seeder pigs were inoculated via the intramuscular route and introduced into contact pigs at ASF pen prevalence of 4 - 5% (n = 20 - 25 pigs) [34]. In this study, the contact pigs as well as the seeder pigs were allowed to chew on the rope during aggregate oral fluid sample collection. ASFV genomic DNA was detected in the whole blood of seeder pigs on 1-3 dpi, in aggregate oral fluids on 3-5 days post contact, 2-3 days prior to the death of the seeder pigs inoculated with virulent ASFV Georgia 2007/1, and 4 - 5 days prior to the death of seeder pigs inoculated with ASFV Malta’78 [34].

In our study, we evaluated pen-based aggregate oral fluid detection of ASFV genomic material using a highly virulent ASFV isolate, obtained from a severe ASF outbreak in Ghana. The seeder pigs were inoculated via the intramuscular route and introduced at a lower ASFV pen prevalence of 3.2% and in contrast to other studies, the seeder pigs did not chew on the rope during aggregate oral fluid sample collection. The seeder pigs developed fever and exhibited clinical signs on 2 - 4 dpi. Disease progression was fast and the pigs succumbed on 5 - 7 dpi, which is consistent with findings in previous studies about faster disease progression in pigs upon intramuscular inoculation with highly virulent strains of ASFV isolates [53]. ASFV genomic DNA was detected in all seeder pigs earlier in the blood on 1 dpi, and later on 2 dpi in the oropharyngeal swabs (Fig. 5A and B).

On the other hand, ASFV genomic material was detected in the aggregate oral fluids as early as 3 dpc in 67% (4/6) of pens and in all 6 pens (100%) on 4 dpc after introduction of the seeder pigs. These findings suggest a 67% chance of detecting ASFV incursion in a naïve swine herd at a comparatively lower ASFV pen prevalence, a scenario that is likely to occur in commercial swine operations. Our study shows that there is 100% chance of detecting ASFV on 4 dpc upon introduction, suggesting that early detection of the virus in a naïve swine herd is possible at the current ASF herd prevalence prior to the onset of apparent clinical signs. In effect, we detected ASFV DNA in aggregate OF on average 3 days prior to the death of seeder pigs, which provides a window for early detection of virus incursion. Clinical signs characteristic of ASF became apparent in some contact pigs on 6-7 dpc, which was 3 - 4 days prior to the occurrence of the first 2 mortality events (Pen A and Pen B, one each) and 6 days prior to significant mortalities (Fig. 9), presenting additional opportunity to initiate timely intervention and disease mitigation measures.

The detection of ASFV genomic material in aggregate oral fluids correlated with the detection of the viral DNA in individual oropharyngeal swabs, as early as 3 - 4 dpc (Table 1). In contrast, detection of ASFV DNA in the blood of individual contact pigs occurred on average 2 days later (1 - 3 dpc), showing earliest detection on 5 dpc and a median positivity of 13.3% (Figure 8D).

These results suggest that aggregate fluid sampling could be a more sensitive and sustainable method for early detection and surveillance of ASF incursion, especially in commercial swine operations, compared to testing individual blood samples. The detection of ASFV in the oral fluids and oropharyngeal swabs earlier than in blood is supported by the pathogenesis of ASFV infection via natural transmission. Following ingestion, which represents the natural route of ASFV transmission in domestic pigs [54, 55], initial replication of the virus occurs in the tonsils and oropharynx [36, 37] and provides plausible explanation for the early detection of the virus in aggregate fluids and oropharyngeal swabs; presenting further evidence of the suitability of aggregate oral fluid sampling as a tool for early detection and surveillance of ASFV. On the other hand, earlier detection in the blood of seeder pigs may be explained by the intramuscular administration, which may allow direct entry of the virus into the circulatory system, replication, and consequently detection early in the blood.

Furthermore, in our study, the contact pigs continue to chew on the rope in significant proportion (> 60%) until 10 dpc (Fig. 8A), even after the appearance of overt clinical signs making it possible to detect the presence of the virus in the oral fluids. Although, the detection of ASFV DNA in aggregate oral fluids remained consistent from 4 -10 dpc across all pens, it occurred at high Ct values suggesting the presence of low amounts of ASFV genomic material. This may pose a challenge to further virus characterization via whole genome sequencing. Or it could be due to environmental contaminants that inhibit PCR sensitivity. Together, these observations suggest the need for research directed at improving the sensitivity (improve Ct values) of aggregate oral fluid detection via improved sample processing and development of enhanced virus capture or enrichment protocols.

Furthermore, the temporal dynamics in the proportion of pigs positive for ASFV genomic material in the blood and oropharyngeal swabs (Fig. 7A and B) and the mortality curves (Fig. 9), is in keeping with previous studies that ASFV in a swine herd spreads slowly but steadily. It may take some time to detect in contact pigs and may be up to few weeks from the beginning of an incursion to the appearance of overt disease [34, 56]. The proportion of pigs positive at the early stages of viral DNA detection varied between pens, which may depend on the hyperactive behavior of the seeder pig and extent of shedding the virus and contaminating the environment. However, with increasing number of pigs getting infected, the virus spread rapidly through the herd and infected nearly 100% of the pigs in all pens by 8 -10 dpc. The uneven spread of the virus among the different pens is in keeping with the lack of uniformity in the mortality curves among the pens (Fig. 9). The first 2 deaths occurred on 10 dpc followed by a significant number of deaths only after almost 2 weeks after introduction of the seeder pigs thus supporting the observation about the stealthy nature of initial ASFV incursion.

A major constraint to enhanced ASFV surveillance in commercial swine operations in North America remains the lack of a reliable non-invasive sampling method for early virus detection. Consequently, we evaluated, as a proof-of-concept, environmental sampling (fecal and water nipple swabs) to support ASF surveillance. Our results show that although early detection of ASFV genomic material in fecal (on 4 dpc) and water nipple (on 2-3 dpc) swabs at pen level may be possible, the lack of consistency at present in detection performance (S3 and S4, Table) indicates the need for further research on optimizing and standardizing the current sampling and extraction protocols.

In our study we generated the seeder pigs via intramuscular inoculation of a highly virulent ASFV. The rapid disease progression in these pigs is in keeping with previous studies [53]. We hypothesize that intramuscular inoculation of the virulent isolate in the current study may have induced significant virus shedding resulting in widespread environmental contamination. It is also hypothesized that disease progression in the seeder pigs and contact pigs as well as environmental contamination could be moderated when seeder or contact pigs are infected through ingestion via contaminated feed, the natural route of ASFV transmission, or if a low or moderately virulent ASFV strain is used to inoculate the seeder pigs. These scenarios may mimic the likely paths for ASFV incursion and manifestation of disease progression in a commercial swine farm in non-endemic settings. Further studies are required to address these hypotheses.

## Conclusion

In conclusion, we have demonstrated that aggregate oral fluid sampling is a sensitive method for early detection of ASFV incursion in naïve swine herds to support disease surveillance in commercial swine operations in North America. Detection of ASFV genomic material in aggregate oral fluids from contact pigs occurred as early as 3 - 4 dpc, 1-3 days earlier than in blood. This correlates with detections in oropharyngeal swabs and further substantiates the utility of aggregate oral fluid sampling as a reliable early detection tool that could be adopted by the swine industry.

## Data availability

All relevant data are in the paper and its supporting information files.

## Competing interests

The authors declare no competing interests.

## Supplementary Materials

Figure S1: Phylogenetic analysis of ASFV isolates based on the *B646L* gene sequences encoding the p72 capsid protein. The tree shows grouping of the Ghana ASFV virus (purple arrow) inoculum with the p72 genotype II cluster. Table S1: Clinical signs and scoring criteria for ASF in experimental pigs; Table S2: ASFV qPCR Ct values of DNA detections in aggregate oral fluids from contact animals in pens at different timepoints post-contact; Table S3: ASFV qPCR Ct values of DNA detections in water nipple swabs from the pens at different timepoints post-contact/dpi; Table S4: ASFV qPCR Ct values of DNA detections in fecal swabs from the pens at different timepoints post-contact/dpi.

## Acknowledgements

The authors would like to thank the Chief Veterinary Officer of Ghana, Dr. Emmanuel Allegye-Cudjoe, and staff of the Accra Veterinary Laboratory of Veterinary Service Directorate, Dr. Joseph Kofi Abuh, Dr. Fenteng Danso, Dr. Patrick Ababio, Daniel Baah, Daniel Arthur, Nana Yaw Tawiah-Yingar Daniel, Dr. Ama Kyerewaa Kusi Appiah and Dr. Eunice Appiah). We also thank Dr. Richard Kwamena Abbiw of the West African Center for Cell Biology of Infectious Pathogen for data compilation, Dr Meyir Ziekah of Wildlife division of Forestry Commission for animal sampling, Doreen Owusu-Ntumy of Animal Research Institute Ghana for laboratory support. We thank Montariel Minor, Catherine Johnson, Andrew Fabian and Amy Berninger at USDA/NVSL/FADDL for laboratory support, and Leslie Blakemore for general support to the study. We thank William Saunders at DHS/PIADC for helping with the shipping arrangements. We thank Lydia Carpenter (APHIS/USDA) and Lynn Wachtman (APHIS/USDA) for their support and contribution. We are thankful to Amy Delgado (CEAH-USDA), Jeffrey Zimmerman (Iowa State University) and Aruna Ambagala (CFIA NCAD, Canada) for sharing their expert knowledge.

## Funding

This study was supported by funding from the United States Department of Agriculture (USDA) Animal and Plant Health Inspection Service (APHIS).

## Author contributions

**Conceptualization:** Suelee Robbe-Austerman, Kathleen O’Hara, Marta Remmenga, Bonto Faburay

**Formal Analysis:** Bonto Faburay, Sherry Johnson, Theophilus Odoom

**Investigation:** Theophilus Odoom, Sherry Johnson, Benita Anderson, William Tasiame, Matilda Ayim-Akonor, Kingsley Kwabena Amoako, Vivian O’Donnell, Lizhe Xu, Corrie Brown, Bonto Faburay

**Methodology:** Bonto Faburay, Suelee Robbe-Austerman, Kathleen O’Hara, Marta Remmenga, Theophilus Odoom, Sherry Johnson

**Project administration:** Bonto Faburay, Corrie Brown, Theophilus Odoom

**Resources:** Suelee Robbe-Austerman, Robin Holland, Bonto Faburay

**Supervision:** Bonto Faburay, Theophilus Odoom

**Visualization:** Bonto Faburay

**Data Curation:** Bonto Faburay, Theophilus Odoom, Sherry Johnson

**Writing - Original Draft Preparation:** Bonto Faburay

**Writing - Review and Editing:** Bonto Faburay, Suelee Robbe-Austerman, Robin Holland, Randall Levings, Vivian O’Donnell, Lizhe Xu, Kate Schumann, Diane Holder, Michelle Zajac, Theophilus Odoom, Sherry Johnson, Corrie Brown

## Notes

### Competing Interest Statement

The authors have declared no competing interest.

