## Supplemental Figure and Tables for "Evaluation of Aggregate Oral Fluid Sampling for Early Detection of African Swine Fever Virus Infection"

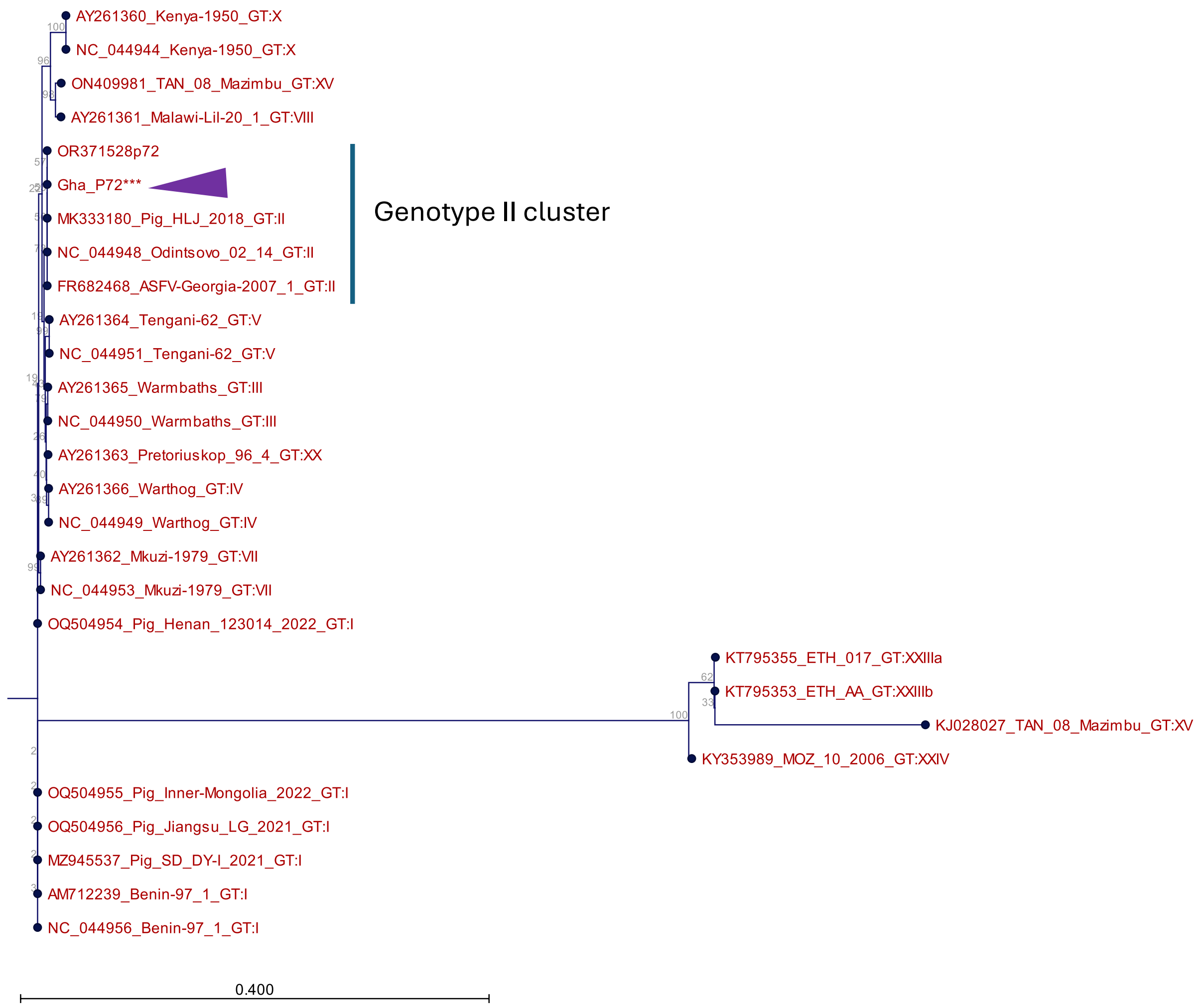


**Supplemental figure 1**. Phylogenetic analysis of ASFV isolates based on the *B646L* gene sequences encoding the p72 capsid protein. The tree shows grouping of the Ghana ASFV virus (purple arrow) inoculum with the p72 genotype II cluster.

**Supplemental table 1.** Clinical signs and scoring criteria for ASF in experimental pigs

| Characteristic | Score | Description |
|---|---|---|
| Behavior and mentation | 0 | Normal, alert response |
| | 1 | Mildly obtunded. Slightly reduced liveliness, stands up unassisted, resists restraint or rectal thermometer |
| | 2 | Obtunded. Reluctant to stand but will do so when assisted; decreased resistance to restraint or rectal thermometer |
| | 3 | Intermittent ataxia, disorientation, can still stand/walk or will not stand/walk even when assisted, still conscious |
| | 4 | Moribund. Non-ambulatory, unconscious/nonresponse |
| Neurologic signs | 0 | Normal |
| | 2 | Unambiguous neurologic signs (e.g. convulsions, seizures) |
| Defecation | 0 | Normal to mildly soft stools |
| | 1 | Profuse watery diarrhea with/without hematochezia or melena |
| | 2 | Severe to marked hematochezia or melena |
| Body temperature | 0 | 38-40 $^{O}$C |
| | 1 | Temperature greater than or equal to 40 $^{O}$C at any point of study |
| | 2 | Temperature greater than or equal to 40 $^{O}$C for at least 2 subsequent days |
| | 3 | Temperature greater than or equal to 41 $^{O}$C |
| | 4 | Temperature less than 38 $^{O}$C |
| | 5 | |

Adapted: Howey et al. [42]; de Carvalho Ferreira et al. [46]

**Supplemental table 2.** ASFV qPCR Ct values of DNA detections in aggregate oral fluids from contact animals in pens at different timepoints post-contact.

| DPC | Aggregate OF qPCR Ct Values | | | | | |
|---|---|---|---|---|---|---|
| | Pen A | Pen B | Pen C | Pen D | Pen E | Pen F |
| 0 | Neg | Neg | Neg | Neg | Neg | Neg |
| 1 | Neg | Neg | Neg | Neg | Neg | Neg |
| 2 | Neg | Neg | Neg | Neg | Neg | Neg |
| 3 | 35.46 | 35.63 | 35.85 | Neg | 36.14 | Neg |
| 4 | 34.63 | 35.54 | 37.55 | 35.7 | 36 | 35.68 |
| 5 | 33.7 | 34.73 | 34.22 | 35.55 | 35.28 | 37.41 |
| 6 | 35.68 | 34.9 | 36.02 | 36.91 | 37.25 | 36.23 |
| 7 | 36.9 | 36.86 | 35.33 | 35.66 | 35.3 | 35.1 |
| 8 | 34.76 | 33.52 | 34.66 | 35.22 | 35.67 | 34.52 |
| 9 | 35.62 | 35.98 | 34.53 | 35.81 | 35.81 | 36.64 |
| 10 | 34.76 | 35.05 | 36.06 | 35.94 | 35.17 | 34.75 |
| 11 | 34.92 | 31.18 | 34.24 | 36.5 | 36.47 | 32.88 |
| 12 | 31.67 | 30.42 | 33.55 | 34.64 | 34.1 | 28.95 |
| 13 | 31.3 | 32.95 | 33.18 | 33.46 | 34.58 | NC |
| 14 | 30.63 | 30.73 | 30.55 | 34.9 | 35.63 | NC |
| 15 | 30.82 | NC | NC | NC | 32.18 | NC |
| 16 | NC | NC | NC | 31.81 | NC | NC |
| 17 | NC | NC | NC | 27.77 | 33.71 | NC |

Note: Red box indicates initial positive detections in aggregate oral fluids in each pen. OF: denotes oral fluid NC: denotes no chewing on the rope due to severity of clinical symptoms.

**Supplemental table 3.** ASFV qPCR Ct values of DNA detections in water nipple swabs from the pens at different timepoints post-contact/dpi.

| DPC/DPI | Water nipple swab qPCR Ct Values | | | | | |
|---|---|---|---|---|---|---|
| | Pen A | Pen B | Pen C | Pen D | Pen E | Pen F |
| 0 | Neg | Neg | Neg | Neg | Neg | Neg |
| 1 | Neg | Neg | Neg | Neg | Neg | Neg |
| 2 | Neg | Neg | Neg | Neg | 36.3 | Neg |
| 3 | 34.5 | 35.1 | 36.9 | 35.5 | Neg | Neg |
| 4 | 33.6 | 36.6 | Neg | Neg | Neg | Neg |
| 5 | Neg | 35.1 | Neg | Neg | Neg | Neg |
| 6 | 35.8 | 33.9 | Neg | 36.8 | Neg | 34.9 |
| 7 | Neg | Neg | Neg | Neg | Neg | 35.4 |
| 8 | Neg | Neg | Neg | 36.6 | Neg | 35.6 |
| 9 | 36.2 | 35.9 | Neg | Neg | Neg | Neg |
| 10 | 33.2 | Neg | Neg | 36.8 | 36 | 36.7 |

Note: Red box indicates initial positive detections in aggregate oral fluids in each pen. Neg denotes negative PCR results.

**Supplemental table 4.** ASFV qPCR Ct values of DNA detections in fecal swabs from the pens at different timepoints post-contact/dpi.

| DPC/DPI | Fecal swab qPCR Ct values | | | | | |
|---|---|---|---|---|---|---|
| | Pen A | Pen B | Pen C | Pen D | Pen E | Pen F |
| 0 | Neg | Neg | Neg | Neg | Neg | Neg |
| 1 | Neg | Neg | Neg | Neg | Neg | Neg |
| 2 | Neg | Neg | Neg | Neg | Neg | Neg |
| 3 | Neg | Neg | Neg | Neg | Neg | Neg |
| 4 | 31.33 | 35.88 | Neg | 33.88 | Neg | Neg |
| 5 | Neg | Neg | Neg | Neg | Neg | Neg |
| 6 | Neg | Neg | 33.74 | 35.18 | Neg | Neg |
| 7 | Neg | 36.06 | Neg | Neg | Neg | 36.65 |
| 8 | Neg | Neg | Neg | Neg | Neg | Neg |
| 9 | 33.45 | Neg | Neg | 35.75 | 34.12 | 32.9 |
| 10 | 29.89 | 35.47 | 31.7 | 34.07 | 34.09 | 31.17 |

Note: Red box indicates initial positive detections in aggregate oral fluids in each pen Neg denotes negative PCR results.
